# The impact of sex differences on perceived pain intensity in pain protocol standardization

**DOI:** 10.1101/2024.12.09.626634

**Authors:** Max Wragan, Niamh O’Connor, Hannah Ashe, Ruairi O’Flaherty, Alice G. Witney

**Affiliations:** Trinity College Institute of Neuroscience, Trinity College Dublin, Ireland; Trinity Centre for Biomedical Engineering, Trinity College Dublin, Ireland; Discipline of Physiology, School of Medicine, Trinity College Dublin, Ireland

**Keywords:** sex differences, heat pain threshold, experimental pain

## Abstract

**Background:** Sex differences have been widely demonstrated in both acute and chronic pain. Sex differences may have wider impact on research design and analysis than already established. This study addresses an important methodological aspect with regards to how sex differences could influence the design of standardized experimental pain protocols used to characterize an individuals’ pain response.

**Methods:** This study addresses sex differences of perceived pain at tonic heat pain threshold (HPT). Participants used a computerized visual analogue scale (CoVAS) to continually rated subjective pain intensity during tonic HPT. Metrics (Mean, Standard Deviation, Maximum) were extracted from the CoVAS to characterize perceived pain.

**Results:** Female participants rated pain intensity at HPT significantly lower than male participants across all extracted metrics used to characterize the coVAS rating. The effect of sex on the mean and standard deviation of pain intensity ratings at HPT was medium, while the effect size of sex on the maximum pain intensity rating at HPT was large.

**Conclusion:** The significant sex differences in perceived pain intensity at HPT indicates that methods of standardization to a specific pain intensity merit sex-specific consideration. Additionally, these observed sex differences underscore the necessity for sex specific design across both pre-clinical and clinical studies of pain.

## Introduction

Pain is a universal phenomenon that can protect against harm, but chronic pain is a disease. Despite being universally pervasive, pain remains difficult to understand because pain is inherently subjective and influenced by a multitude of biological and social factors [1]. Biological sex is an established influencer of pain perception. Prevailing research indicates females demonstrate higher sensory and nociceptive discrimination and detection relative to males, attributable to biological sex differences across molecular, cellular, and system levels[1-3]. Sex differences occur in both chronic pain and experimental pain studies[4]. The importance of sex differences in pre-clinical and clinical research is gaining increasing attention [5, 6] with the need to design and analyze data with the potential impact of sex differences carefully considered[6].

Chronic pain acknowledgment as a disease by the international classification of diseases (ICD-11), increases the need for objective assessment tools. The mechanisms underlying chronic pain are poorly understood, but some patients are thought to have defective endogenous descending pain pathways[7]. Consequently, experimental pain protocols have been developed to assess descending modulatory pain pathways, including offset analgesia (OA) and conditioned pain modulation (CPM). These experimental pain protocols feature individualized pain thresholds to standardize these protocols against subjective differences in pain [8-10]. A typically used metric is heat pain threshold (HPT), the lowest temperature at which an individual experiences pain. HPTs range from 42-49°C, and although sex differences are thought to be larger to mechanically induced pain relative to noxious heat[11], women typically report HPTs a single or few degrees lower than men [1, 12-14]. Further, HPTs are regularly used to standardize thermal stimuli within experimental pain protocols of temporal summation, OA, and CPM. However, pain intensity ratings of individuals’ HPTs are presently insufficiently studied and understood, despite considerable research demonstrating the sex-based differences in temperatures of HPT[1, 12, 13].

The ambiguity surrounding sex-based differences in pain intensity at HPT sources from a lack of targeted research and precise tools measuring pain intensity. Previous findings are complicated by the different pain measurement tools used across studies, including visual analogue scale (VAS) and numerical rating score (NRS) ratings that often are collected just once post-procedure[12, 15].Inconsistent sex differences have been found in the perceived intensity of pain at a given noxious temperature stimuli, but it remains unclear if sex-based differences extend to the perceived pain intensity of heat pain thresholds[1, 16-18].

To resolve the limitations identified in previous pain measurement tools, the TSA-II (Medoc, Israel) delivers computer-controlled pain and measures pain intensity in real time using a computerized visual analogue scale (CoVAS). One study, Wise et al. 2002, included CoVAS to compare sex differences in pain intensity; however, the analysis was restricted to a single instantaneous metric[19]. Given the current importance of HPTs in standardizing experimental pain protocols, correct standardization is essential. This study examines this methodological gap to ensure optimal standardization of experimental pain protocols by evaluating sex-based differences in perceived pain intensity at HPT.

## Material and methods

### Participants and Experimental Ethics

The study was a retrospective data analysis of sex differences in pain ratings in a dataset of ramp and hold heat pain protocols. Therefore, there was double blinding during data collection. 39 participants, 16 females and 23 males were used in the study. Participants had a mean age of 22.45 years old.

Data regarding participants perceived pain intensity at HPT was collected twice during two separate sessions performed a week apart for all participants.

As part of the experiment’s recruitment process for the ramp and hold trials, participant eligibility was screened via a medical questionnaire for the following exclusion criteria: epilepsy, chronic pain, migraines, motor diseases, cochlear implants, neurological issues, and psychiatric conditions. Upon meeting inclusion criteria, all participants provided their written informed consent. Ethical approval for the data collection was granted by the Faculty of Health Sciences, Trinity College Dublin, Ireland. All experimentation was performed in accordance with institutional requirements and the Declaration of Helsinki (2013).

### Painful Thermal Stimulation

Pain was induced via thermal heat stimuli using a TSA-II Neurosensory Analyzer. Heat was applied via the 16×16 mm thermode Peltier TSA-II attachment with a Velcro ® strap to secure the thermode to the right forearm of participants. To help ensure data accuracy, 60s of rest and minor thermode repositioning followed any repeat measures. Importantly, minor thermode repositioning reduces concerns of sensitization without altering pain response sensitivity [20].

### CoVAS Pain Measurement

Participants’ subjective pain intensity was measured and recorded with a computerized visual analogue scale (CoVAS, Medoc Ltd, Israel). CoVAS is an industry gold standard tool that allows participants to continually quantify their pain levels by displacing the marker of a 10 cm slider between the two endpoints of ‘no pain’ and ‘most intense pain imaginable’. The level of displacement from the ends corresponds to a numerical whole number between 0 and 100. The advantage of CoVAS measurement over other pain measurement tools is the data collection rate of 9 Hz. This frequency of data collection captures continual real-time dynamic aspects of pain ratings that are overlooked by other methods of pain evaluation. Prior to data collection, participants trained with the CoVAS equipment, reducing non-significant data variation to ensure data reflected participants’ sensations of pain instead of temperature [20-22].

### Identifying Heat Pain Thresholds

Prior to any CoVAS measurements, HPTs were individually established for all participants [8, 10]. Participants’ HPTs were determined by increasing the temperature of the thermode at a rate of 1.5°C/s from baseline until the temperature at which participants first reported experiencing pain. This process was performed three times, punctuated by 10s intervals of rest between HPT trials. If all three HPTs varied by less than 0.8°C, the three values were averaged to serve as the participants’ HPT. However, if any of the three recorded pain thresholds varied by more than 0.8°C, HPT protocols were repeated until the variation reduced to less than 0.8°C, such that subsequent protocols could rely on a mean HPT with high precision.

### Pain Intensity at Heat Pain Thresholds

Ramp and hold protocols were used to characterize participants’ pain response during HPT exposure (Figure 1) [23]. Ramp and hold trials are frequently used as control trials within pain clinical research and adjacent fields [24-27]. During ramp and hold trials, participants’ stimulating thermode was increased from baseline temperature to individuals’ HPT at a rate of 1.5°C/s. Once HPT was reached, this temperature was held for 30s before the thermode was returned to baseline temperature at a rate of 6°C/s. Throughout this entire procedure, participants’ subjective sensation of pain was captured in real-time using the CoVAS.

**Figure 1.**
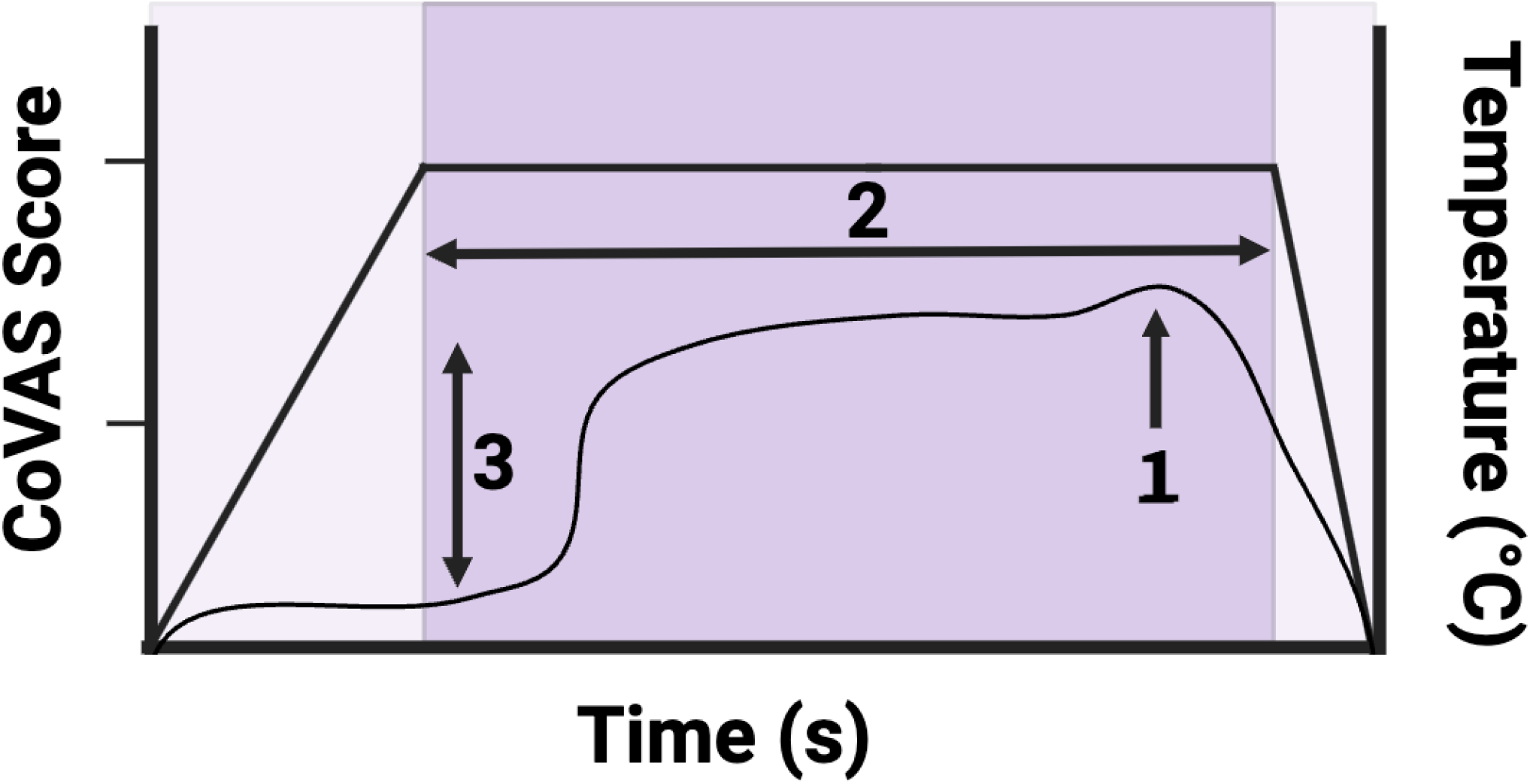
Ramp and Hold Control Trial Profile. The figure visualizes the ramp and hold control trial temperature protocol with the black line, where thermal stimulus is increased from baseline to participants HPT at a rate of 1.5°C/s. Once HPT is reached, the HPT is held for 30 s while participants use the CoVAS to continually rate their pain sensation in real time. At the end of 30 s period, the temperature returns to baseline at a speed of 6°C/s. Additionally, extracted variables are visualized with 1 = Maximum VAS; 2 = Mean VAS; 3 = Standard Deviation VAS.

## Statistical Methods

### Extraction of Metrics

The measurements extracted from participants’ CoVAS profile for analysis were selected to capture temporal and magnitude characteristics of participants’ pain profiles. A custom written MATLAB script extracted the metrics (Figure 1)(Version 9.0, The MathWorks Inc., MA, USA). This experiment extracted the mean and maximum VAS to capture response magnitude, and standard deviation of VAS score recorded during the 30s HPT plateau phase was used to capture response variance. Ramp and hold HPT experiments are relatively simple, thus those three variables were determined sufficient to characterize participant response (Harris et al., 2018).

#### Definitions of Ramp and Hold Trial Extracted Values

1. Maximum – Magnitude of highest pain score recorded during HPT plateau
2. Mean – Average VAS score during HPT plateau
3. Standard Deviation – Standard deviation of VAS score during HPT plateau

### Statistical analysis

Statistical analysis was performed in SPSS (IBM SPSS Statistics, Version 27; IBM Corp), with data visualization performed within PRISM (GraphPad Prism, Version 9; GraphPad Software Inc).

First, the two time points for each participant were averaged to reduce any single data collection variation per participant. Using participants’ averaged data, descriptive statistics were collected for all variables, including mean and standard error of the mean. Normality tests, histograms, Q-Q plots, and boxplots were performed for all variables and independent sample t-tests were performed to compare female and male participants for all variables. This evaluation for sex-based differences was supplemented with scatterplots visualizing data distribution. Effect size calculations were performed to provide a standardized measure of magnitude for the significant effect of sex identified on the perception of pain, as measured by mean, maximum, and standard deviation of VAS scores [19]. Effect size was determined by subtracting the average female VAS from the average male VAS and then divided by the data’s pooled standard deviation.

## Results

### Assessment of VAS responses during HPT plateau

Descriptive statistics characterizing the mean, standard error of the mean, and range of the data when separated by sex is summarized in Table 1. Irrespective of sex the average mean VAS was 31.96, the average mean maximum was 41.91, and the average standard deviation was 8.15. However, when separated by sex, female participant data averaged a lower mean (23.70 cf 37.71), maximum (32.53 cf 48.43), and standard deviation VAS (6.17 cf 9.53) score from the thirty second HPT plateau relative to male participants.

**Table 1.** Ramp And Hold Descriptive Statistics. The table displays descriptive statistics for the mean VAS, maximum VAS, and standard deviation VAS averaged across the two test dates differentiated with respect to sex.

|  | Mean | S.E.M. | Maximum | Minimum |
| --- | --- | --- | --- | --- |
| Female Mean VAS | 23.70 | 3.95 | 60.77 | 4.95 |
| Male Mean VAS | 37.71 | 4.02 | 89.69 | 10.14 |
| Female Maximum VAS | 32.53 | 4.06 | 64.50 | 9.00 |
| Male Maximum VAS | 48.43 | 4.55 | 95.50 | 14.50 |
| Female Standard Deviation VAS | 6.17 | 0.99 | 13.17 | 1.95 |
| Male Standard Deviation VAS | 9.53 | 1.06 | 22.47 | 2.22 |

### Evaluating sex differences

The analysis revealed significant differences between females and males pain response to a ramp and hold heat pain stimulus. The t-tests for sex-based differences identified a significant effect of sex on mean VAS (p = 0.022), maximum VAS (p = 0.018), and standard deviation VAS (p = 0.033). Thus, the mean, maximum, and standard deviation VAS scores reported by females during the HPT plateau were significantly lower than male VAS metrics. The scatter plots in figure 2 visualize the data and sex-based differences.

**Figure 2.**
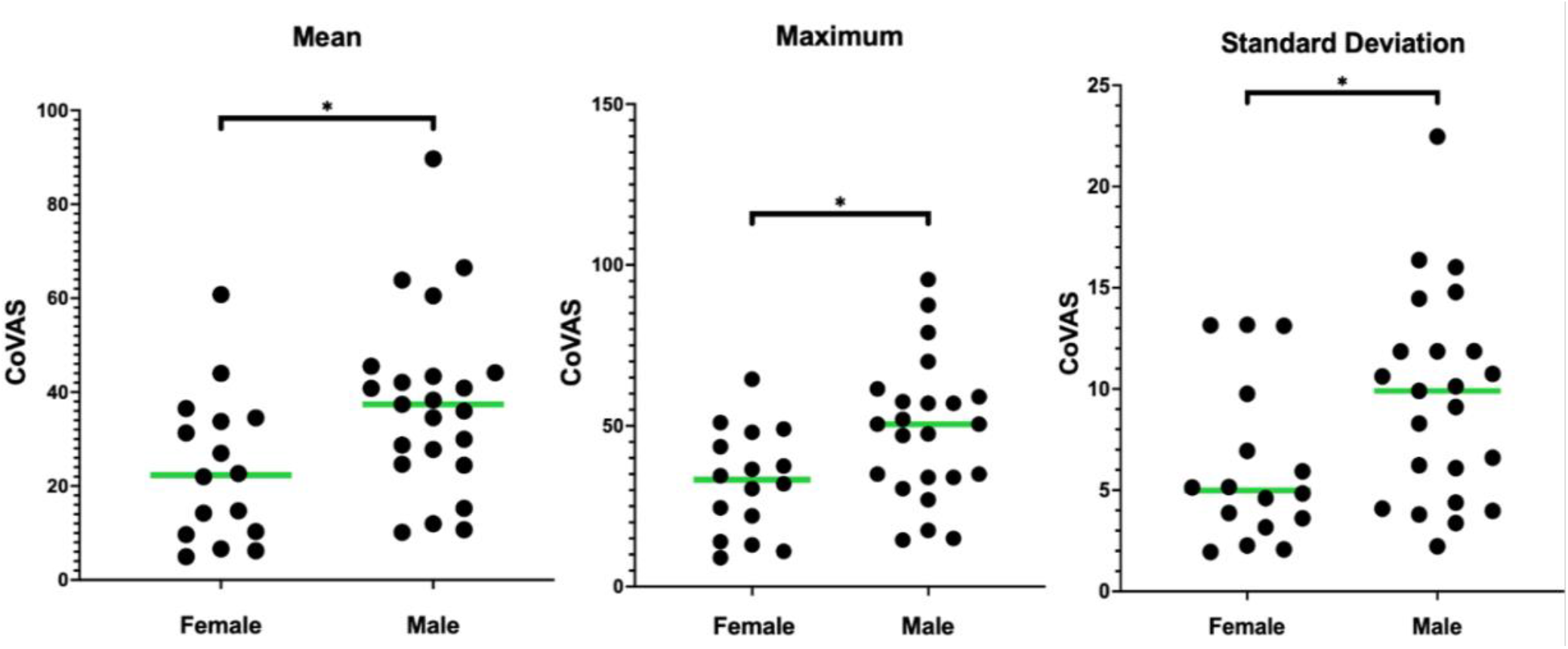
Scatter Plot of Extracted Variables Separated by Sex. The scatter plots visualize the sex-based differences that were revealed via t-tests. Sex has a significant effect on all three ramp and hold variables: mean, maximum, and standard deviation.

### Quantifying the effect of sex on pain perception

The effect size of sex on the mean VAS was medium boarding large (d = 0.795). The effect of sex on the maximum VAS was large (d = 0.827). The effect of sex on standard deviation of VAS was medium (d = 0.734)

## Discussion

Sex differences in pain perception are well established. Recently, the importance of consideration of the impact of sex differences at all stages of study design and analysis has been emphasized [4, 6]. There is limited research investigating the relation between sex and the perception of pain at HPT, a metric regularly integrated into the experimental design of numerous studies across the pain research field. Although HPT attempts to standardize pain experimentation through individualization, the lack of robust research regarding potential sex differences in the perceived pain intensity at HPT threatens to undercut experimental standardization efforts. Therefore, this study addresses this technical consideration in pain standardization to close a foundational gap that should be considered in the standardization of pain protocols. The identification of the significant medium to large effect of sex on pain perception at HPT by this study further underscores the need to account for sex-based differences across pain research.

Within previous clinical pain research, both acute and chronic pain models show sex differences. Our findings on sex differences in pain perception may have different implications for acute and chronic pain. Acute pain focuses on immediate sensory responses and could be impacting by the likely underlying mechanisms endogenous pain system sex-differences responsible for the sex-based discrepancy in our results; however, sex-differences in endogenous pain systems and their corresponding impact on chronic pain modulation is likely more complex and sex-specific, considering the greater complexity and the established greater prevalence of chronic pain in women[28].

Previous research has demonstrated females have lower temperature heat pain thresholds, as females are more sensitive and responsive to nociceptive stimulus[1, 2, 29]. Previous research also evidenced that women frequently report higher subjective pain intensity ratings relative to men across many diagnostic and experimental contexts[28]. However, this study reveals that even when participants are standardized to their individual threshold temperature, the temperature at which they first start to experience pain, women regard this threshold sensation of pain as less painful. This identified sex difference in pain perception at threshold introduces complexity into attempts at individually standardizing research protocols, particularly those examining descending pain modulation. Individualizing these protocols to accommodate the heightened pain sensitivity of women may not resolve sex differences in subjective pain perceptions of threshold stimuli.

Despite the research of this study and previous research identifying sex-based differences in pain, the underlying systems contributing to these well documented sex differences remain unclear. Numerous studies have identified female sensory and nociceptive systems are more sensitive than males, as evidenced by the frequently lower pain threshold values identified in female participants, but the biological systems implicated underlying these differences are complex and interacting[1, 28]. The same systems responsible for the sex differences in threshold temperature and perception intensity at fixed temperatures are likely to underlie the sex-based differences of perceived pain at HPT.

Research has identified female A-delta afferents are more responsive to sensitization than males, representing a serious possible contribution towards the sex-based differences in perceived pain at HPT[29]. In previous research, these sex-based differences in A-delta afferents were indicated to underlie women’s higher pain reports during early phase noxious stimulus. However, interestingly women also have demonstrated greater attenuation to painful stimuli, indicating women are more acutely sensitive and responsive to pain. In addition to these nociceptive fibers, sex differences are also well documented at the level of nociceptive structures and systems. Studies of acute and chronic pain models have demonstrated higher analgesic activity of male higher nociceptive structures like the PAG and RVM relative to females, attributed to reduced potency of mu agonists in females and sexually dimorphic organization and functional of the PAG-RVM circuit [30].

In addition to these system level sex differences, cellular and molecular differences are also attributed to underlie sex differences in pain. Research on the TRPV1 channel, responsible for noxious stimuli sensation including noxious heat, has demonstrated women experience greater pain sensation from TRPV1 agonist exposure relative to men [31]. TRP channels have also shown modulation in response to sex hormones [31]. Research also indicates that microglia play a prominent role in male pain perception mechanisms, but not in females; instead, t-cells are more centrally involved in female pain perception mechanisms [32]. Sexually dimorphic pathways of pain processing and response could underlie a multitude of sex differences in pain experiences between women and men. Further, hormonal differences are implicated to alter pain response across sex. In the case of microglia, research has identified testosterone stimulates and controls microglia dependent pain pathways [32]. Other research has identified estrogens increase NMDA receptor excitability, thereby enhancing nociceptor excitability in females[33]. This enhanced excitability of nociceptors by a major female sex hormone provides a reasonable explanation to the greater sensitivity of female participants to noxious heat stimuli than males. Research on these cellular and molecular sex differences is largely restricted to animal model research, but these systems are highly conserved and indicating these sex differences in pain perception parallels humans[34].

The role of menstrual cycle phases and hormone fluctuations in female pain perception was not explicitly evaluated for this study, as our sex-difference findings in pain perception were yielded as a retrospective analysis of data collected on HPTs. However, given the role of hormones in nociceptive and sensory perception, future research with specific consideration towards hormonal fluctuations would build on the presented findings and refine our understanding on the biological contributions to sex-different pain modulation in clinical and experimental settings. Additionally, although gender and social factors were not an explicit focus of this research, societal roles and cultural influences are frequently investigated in relation to pain perception, with many studies identifying various influences of gender identification and pain[1]. Future studies would benefit from specific consideration that individualizes pain metrics’ interaction with gender through questionnaires like the gender role expectations of pain questionnaire [25, 35].

Ultimately, the findings of this study advocate for isolated consideration of pain metrics across sex and gender. The findings of this study also introduce additional complexity into all pain research attempting to standardize pain perception using individualization, raising questions of how to best standardize pain research and medical procedures. Future research is needed to determine if thresholds as they are currently defined as the first level of stimulus at which individuals experience pain are sufficient, or if pain research would be better served by standardizing to a specific subjective pain rating value to better standardize pain research to the inherently subjective nature of pain.

## Conclusion

Experimental pain protocols are being developed to attempt to stratify patient groups based on their pain report to dynamic pain stimuli, as well as the evaluating sensory threshold changes using quantitative sensory testing. These pain protocols, for instance conditioned pain modulation (CPM) are typically individualized, often to a participant’s HPT. However, sex differences in the pain perception of individualized pain variables are insufficiently understood and may mean that these pain protocols do not adequately account for such differences. This experiment considers the effect of sex on pain intensity perception at heat pain threshold using CoVAS to capture a comprehensive and detailed pain perception profile. Statistical analysis across all extracted measurements from participants’ pain profiles revealed a significant effect of sex, with effect sizes that ranged medium to large highlighting and further characterizing differences in male and female nociception processes. Future research should consider pain perception of female and male participants separately to continue characterizing the complicated interaction between sex and pain. Additionally, efforts to standardize pain research against individualized and subjective aspects of pain necessitates further consideration to determine the most effective way to standardize pain protocols and research while adequately accounting for the complex interaction between sex and pain.

## Acknowledgments

Dr. Cathal Walsh (Insight Statistical Consulting) provided statistical expertise for the project.

## Credits

Conceptualization (MW), Methodology (NOC,HA,ROF,MW,AW), Data collection (NOC, HA, ROF), Data collation and metric extraction (MW), Statistical analysis (MW), writing of first draft (MW), review and editing (MW, AW).

